# Technology-aided assessment of functionally relevant sensorimotor impairments in arm and hand of post-stroke individuals

**DOI:** 10.1101/2020.04.16.044719

**Authors:** Christoph M. Kanzler, Anne Schwarz, Jeremia P.O. Held, Andreas R. Luft, Roger Gassert, Olivier Lambercy

## Abstract

**Background:** Assessing arm and hand sensorimotor impairments that are functionally relevant is essential to optimize the impact of neurorehabilitation interventions. Technology-aided assessments should provide a sensitive and objective characterization of upper limb impairments, but often provide arm weight support and neglect the importance of the hand, thereby questioning their functional relevance. The Virtual Peg Insertion Test (VPIT) addresses these limitations by quantifying arm movements and grip forces during a goal-directed manipulation task without arm weight support. The aim of this work was to evaluate the potential and robustness of the VPIT metrics to inform on sensorimotor impairments in arm and hand, and especially identify the functional relevance of the detected impairments.

**Methods:** Arm and hand sensorimotor impairments were systematically characterized in 30 chronic stroke patients using conventional clinical scales and the VPIT. For the latter, ten previously established kinematic and kinetic core metrics were extracted and compared to conventional clinical scales of impairment and activity limitations. Additionally, the robustness of the VPIT metrics was investigated by analyzing their clinimetric properties (test-retest reliability, measurement error, and learning effects).

**Results:** Twenty-three of the participants, the ones with mild to moderate sensorimotor impairments and without strong cognitive deficits, were able to successfully complete the VPIT protocol (duration 16.6 min). The VPIT metrics detected impairments in arm and hand in 90.0% of the participants, and were sensitive to increased muscle tone and pathological joint coupling. Most importantly, moderate to high significant correlations between conventional scales of activity limitations and the VPIT metrics were found, thereby indicating their functional relevance when grasping and transporting lightweight objects as well as dexterous finger manipulations. Lastly, the robustness of three out of the ten VPIT core metrics in post-stroke individuals was confirmed.

**Conclusions:** This work provides evidence that technology-aided assessments requiring goal-directed manipulations without arm weight support can provide an objective, robust, and clinically feasible way to assess functionally relevant sensorimotor impairments in arm and hand in chronic post-stroke individuals with mild to moderate deficits. This allows better identifying impairments with high functional relevance and can contribute to optimizing the functional benefits of neurorehabilitation interventions.

Retrospectively registered: clinicaltrials.gov/ct2/show/NCT03135093

## 1 Introduction

Stroke is a leading cause of acquired adult disability [1]. The incident commonly causes chronic sensorimotor deficits in arm and hand (impairments) [2, 3]. Impairments that are functionally relevant are especially critical for a ected individuals, as these impairments reduce the spectrum of activities that an individual can perform (activity limitations) and determine the level of dependence on caregivers. Neurorehabilitation attempts to decrease the level of disability through physical therapy [4, 5]. Achieving successful rehabilitation, with clear benefits for the independence of individuals, typically requires the identification and therapy of functionally relevant impairments [6–8].

Conventional clinical scales are the standard to evaluate upper limb sensorimotor impairments in research studies and the described impairments mostly show strong links to activity limitations (i.e., functional relevance) [9–13]. However, conventional assessments commonly rely on subjectively rated ordinal scales with ceiling effects that are not sensitive enough to detect fine changes in impairments and even introduce bias when attempting to model sensorimotor recovery [14–16]. Hence, providing a more fine-grained and objective assessment of functionally relevant sensorimotor impairments with sensitive scales should be of primary interest to neurorehabilitation researchers.

Digital health metrics extracted from technology-aided assessments can provide objective and traceable descriptions of upper limb behaviour on sensitive, continuous scales without ceiling effects [17–19]. However, the majority of instrumented assessments focuses on characterizing impairments during isolated planar joint movements while supporting the arm against gravity [20–23]. This neglects the importance of hand impairments and shadows the effect of certain deficits, such as weakness [19], which are both fundamental when performing daily activities. This questions the functional relevance of these assessments.

More recently, technology-aided approaches started emphasizing the importance of assessing impairments during tasks involving arm movements and hand manipulations, without providing arm weight support [24–27]. Such tasks are expected to provide crucial information on fine upper limb impairments in individuals with mild to moderate disability levels and are promising to better identify functionally relevant impairments. However, existing approaches typically rely on time-consuming and complex measurement setups that reduces their clinical applicability. Further, they mostly focus on kinematic metrics and do not quantify grip force control and its essential role in daily life activities. Also, the clinimetric properties of such digital health metrics are often insu ciently evaluated, thereby challenging their interpretability and acceptability as clinical endpoints [17, 28].

The primary objective of this work was to evaluate the potential of digital health metrics, extracted from the Virtual Peg Insertion Test (VPIT) in chronic post-stroke individuals, to inform on arm and hand sensorimotor impairments, and especially characterize the functional relevance of the detected impairments. The VPIT addresses the limitations of existing technology-aided assessments by recording movement and grip force patterns during a virtual goal-directed manipulation task requiring coordinated arm and hand movements [29–33]. Previous research indicated the feasibility of the approach in neurologic individuals with mild to moderate sensorimotor impairments. In addition, ten digital health metrics capturing sensorimotor impairments have been established for the VPIT and allowed accurately discriminating neurologically intact and a ected individuals [33]. However, other clinimetric properties (reliability, measurement error, learning effects) have only been evaluated in una ected subjects. Hence, the secondary objective of this work was to characterize the clinimetric properties of the VPIT metrics in chronic post-stroke subjects and ensure their pathophysiological interpretation and robustness.

To achieve these objectives, we strived 1) to systematically characterize arm and hand sensorimotor impairments in 30 chronic stroke subjects using the digital health metrics of the VPIT and conventional scales. In addition, we aimed 2) to characterize the functional relevance of the detected impairments by correlating them to conventional assessments of activity limitations. Lastly, we intended to 3) analyze test-retest reliability, measurement error, learning effects, and concurrent validity of the VPIT metrics. We hypothesized that the technology-aided assessment with the VPIT provides fine-grained and robust information about sensorimotor impairments in arm and hand that are functionally relevant. This is expected to lead to high correlations between the digital health metrics of sensorimotor impairments and conventional scales of activity limitations. This work contributes to better linking the technology-aided assessment of impairments with activity limitations, thereby opening new avenues to optimize the benefits of neurorehabilitation interventions by identifying functionally relevant therapy targets.

## 2 Methods

### 2.1 Virtual Peg Insertion Test (VPIT)

The VPIT as an upper limb sensorimotor assessment has been described in detail in previous work [29, 30, 33]. In short, it consists of a commercial haptic end-e ector device (PhantomOmni or Geomagic Touch, 3D Systems, USA), a rapid-prototyped grasping force sensing handle, and a virtual reality environment on a personal computer (total material costs approximately 4000 USD). The virtual reality environment displays a virtual pegboard task that requires the insertion of nine virtual pegs into nine holes. More specifically, a virtual cursor can be controlled through the coordination of end-e ector movements and applied grasping force. To pick up a peg, the cursor first needs to be spatially aligned with the peg. Subsequently, a grasping force of at least 2N has to be maintained to transport the peg towards a hole. The peg can be released in a hole upon a reduction of the grasping force below 2N.

Recently, a processing pipeline has been defined to extract and normalize ten kinematic and kinetic digital health metrics from VPIT data (position and grip force sampled at 1 kHz, details in [33]). For this purpose, data is low-pass filtered and temporally segmented into the *transport* (gross movement from peg pickup until insertion), *return* (gross movement from peg insertion to next pickup), and *peg approach* (fine movement after return and before transport), *hole approach* (fine movement after transport and before return). Subsequently, metrics were defined for each of these confined phases to quantify di erent aspects of upper limb sensorimotor impairments.

Smooth movements, represented through a bell-shaped velocity profile, are a hallmark of intact motor control [34]. Movement smoothness was quantified using the normalized logarithmic jerk metric (*log jerk*) calculated during *transport* and *return* as well as the spectral arc length metric of the velocity signal during return (*SPARC return*) [35–37]. Similarly, ballistic movements of una ected individuals are effcient and tend to follow a trajectory close to the shortest path between start and target. Movement efficiency was characterized using the *path length ratio* (shortest possible distance divided by the actually covered distance) during *transport* and *return* [38]. Movement speed was quantified using the maximum velocity during *return* (*velocity max. return*) and the endpoint-precision of the ballistic movement using the jerk metric calculated during the peg approach (*jerk peg approach*). Further, three metrics describing the smoothness of grip force coordination during di erent movement phases were defined. This included the number of peaks in the grip force rate (first time-derivative of grip force) during *transport* (*grip force rate num. peaks transport*). Additionally, the SPARC was applied to grip force rate data recorded during *transport* (*grip force rate SPARC transport*) and *hole approach* (*grip force rate SPARC hole approach*). The clinimetric properties (test-retest reliability, measurement error, learning effects) of all ten metrics have been positively evaluated in neurologically intact subjects [33]. In addition, all metrics indicated strong discriminative ability between a normative reference population and a group of 89 neurologically a ected subjects, thereby demonstrating their ability to capture sensorimotor impairments.

For all metrics, mixed effect models were generated to compensate for confounding factors such as age, gender, tested body side, and whether the test was performed with the dominant body side or not. Further, the value of each metric was normalized with respect to the median and variability of a reference population containing 120 unimpaired subjects (age 20-80 years, 60 female) that performed the VPIT. Lastly, each metric was additionally normalized with respect to the neurologically a ected subject in the VPIT database that showed worst performance in a specific metric. This resulted in metrics being defined on an unbounded scale, theoretically ranging from]*−1*%, +*1*%[, with 0% indicating median task performance of the reference population and 100% worst recorded task performance [33].

### 2.2 Conventional clinical assessments

A battery of conventional clinical assessments was performed to capture the heterogeneity of sensorimotor impairments and activity limitations.

#### Sensorimotor impairments

Hand and wrist impairments as well as flexor/extensor synergies in shoulder, elbow, wrist, and hand were described using the Fugl-Meyer assessment for the upper extremity (FMA-UE) [14]. It focuses especially on abnormal muscle activation patterns that prohibit isolated joint movement of shoulder, elbow, wrist, and hand. The assessment requires the subject to perform specific movements that are known to elicit this coupling, which are subjectively scored on a ordinal scale (0: cannot perform, 1: performs partially, 2: performs fully), leading to a ceiling effect at score 66. The assessment takes approximately 30 minutes to administer [14, 39].

Cognitive impairments were rated with the Montreal cognitive assessment (MOCA), which consists of simple tasks such as drawing, object naming, memory recall, reading, and mathematical operations (0: worst score, 30: best score) [40].

Resistance against passive movements due to increased muscle tone (referred to as spasticity) in shoulder internal rotators, biceps, triceps, wrist flexors and extensors, as well as finger flexors and extensors were defined with the Modified Ashworth Scale (MAS) that involves the passive movement of the respective joint [41]. Trained clinical personnel performed and rated each movement subjectively (0 normal tone, 5 rigid), which takes in total up to 5 minutes time [39]. The ratings were combined into a single score describing overall upper limb muscle tone with a ceiling effect at value 35.

Somatosensory impairments of upper arm, lower arm, hand, and finger was measured based on the Erasmus modified Nottingham sensory assessment (EmNSA) that focuses especially on tactile sensation, sharp-blunt discrimination, two-point discrimination, and proprioception [42]. Therein, the skin was stimulated with di erent objects and the subject had to define touch modality (e.g., light touch vs pressure) or location. Further, proprioception was evaluated by passively moving the participants joints, by asking the subject to indicate the perceived direction of movement, and by comparing the indicated with the actual direction. Each task was scored from zero (no proprioception) to two (normal), leading to a total combined upper limb score of maximal 40 points. The evaluation takes approximately 10-15 minutes to administer [42].

#### Activity limitations

The ability to coordinate precise object manipulations with gross arm movements was evaluated with the Action Research Arm Test (ARAT), which requires the transfer of small and large items with multiple handgrip types from the bottom to the top of a shelf [43, 44]. Each subtask was subjectively rated from zero (task not possible) to three (normal task performance), leading to a maximal possible performance of 57 points.

Fine manual dexterity was evaluated with the time to insert nine small physical pegs into nine corresponding holes without requiring active lifting of the arm against gravity, as defined by the Nine Hole Peg Test (NHPT) [45, 46].

Lastly, gross manual dexterity was reported through the Box and Block Test (BBT), which requires the transport of as many blocks as possible within one minute across a physical barrier while actively lifting the arm against gravity [44, 47]. For the BBT and NHPT, the outcome measure was normalized with respect to the publicly available reference data to account for the influence of age, gender, and tested body side.

### 2.3 Participants and procedures

Thirty post-stroke subjects were recruited at the University Hospital of Zurich (Zurich, Switzerland) and the cereneo, Center for Neurology and Rehabilitation (Vitznau, Switzerland) as part of an observational study (ClinicalTrials.gov Identifier: NCT03135093) that used the VPIT as a secondary outcome next to a battery of clinical assessments focusing on sensorimotor impairments (FMAUE, MOCA, MAS, EmNSA). The VPIT protocol consisted of receiving standardized instructions, familiarizing with the task by inserting all nine pegs once (data not analyzed), and subsequently performing five repetitions (i.e., inserting all nine pegs five times). The protocol was performed with the most a ected and less a ected body side, given that both of them might be a ected by sensorimotor impairments [48]. Further, the subjects were enrolled into a second measurement session including a repetition of the VPIT protocol and further clinical assessments focusing on activity limitations (BBT, NHPT, ARAT).

All participants gave written informed consent, and all procedures were approved by the local Ethical Committees (ID 2016-02075 and BASEC:2017-00398). Recruited were subjects of at least 18 years age with chronic (i.e., at least 6 month ago) ischemic stroke with at least partial ability to lift the arm against gravity and flex and extend the fingers. Exclusion criteria were other concomitant diseases a ecting the upper limb, severe sensory deficits, and severely increased muscle tone that considerably limits range of motion.

Participants started the VPIT assessment with the most a ected body side and were instructed to perform the task as fast and precise as possible. The starting position was approximately 45° shoulder abduction, 10° shoulder flexion, and 90° elbow flexion. Subjects received live feedback about the duration of each VPIT repetition through a timer displayed on the computer screen.

### 2.4 Data analysis

#### Characterization of upper limb sensorimotor impairments and activity limitations

The presence of upper limb impairments was quantified using the ten VPIT metrics and conventional scales. For the VPIT, previously established cut-o s based on the 95*^th^*-percentile of the normative reference population were used to define individuals with abnormal behavior (binary value) in each metric. Afterwards, one value per factor (i.e., physiological constructs previously identified through an explanatory factor analysis) was generated by pooling the information about the presence of abnormal behaviour across all metrics within this factor via the maximum (i.e., factor indicated as abnormal if at least one metric within this factor was abnormal). For the NHPT and BBT, abnormal behaviour was defined if task performance was worse than 1.96 times the standard deviation (corresponding to 95*^th^*-percentile) of the publicly available normative reference population [45, 46]. According to the ARAT, activity limitations were present if the score was below 55 [13]. All other conventional scales indicated the presence of impairments if the full score was not reached.

#### Correlation of upper limb sensorimotor impairments with activity limitations

To analyze how both VPIT metrics and conventional impairment scales relate to conventional assessments of activity limitations, Spearman correlation coe cients (*ρ*) were calculated. For the correlation analysis, only data from the most a ected side (*ρ_ma_*) and the first testing session was included to avoid the influence of ceiling effects in the conventional scales for the less a ected body side and learning effects across sessions, respectively. Bonferroni correction was applied for each tested hypothesis to account for multiple comparisons. The intervals suggested by Hinkle et al. were used for interpreting the correlation coe cients: very high: *ρ_ma_* ≥ 0.9; high: 0.7 *ρ_ma_ <*0.9; moderate: 0.5 *ρ_ma_ <*0.7; low: 0.3 *ρ_ma_ <*0.5; very low: *ρ_ma_ <*0.3 [49].

#### Test-retest reliability, measurement error, learning effects, and concurrent validity of VPIT metrics

The evaluation of the clinimetric properties was guided through a previously defined framework for the selection and validation of digital health metrics [33]. More specifically, the repeatability of the VPIT metrics was quantified by their ability to discriminate di erent subjects across measurement sessions (test-retest reliability) and the measurement error of the task and assessment platform [33, 50, 51]. The former was defined using the intra-class correlation coe cient (ICC A,k). Metrics with an ICC*>*0.7 passed the evaluation. The latter was characterized using the smallest real di erence (SRD), which defines a range of values for that the assessment cannot distinguish between measurement noise and an actual change in the underlying physiological construct. The SRD was defined as 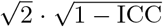 [52, 53]. The SRD was further normalized (SRD%) with respect to the range of observed values of a metric to enable a comparison across metrics. A cut-o of SRD 30.3% was applied to define metrics that have highest potential to sensitively measure sensorimotor recovery [33]. As the smallest real di erence and thereby the responsiveness of a metric strongly depends on the intra-subject variability, the standard deviation across all repetitions of the VPIT was visualized. In addition, Bland-Altman plots were constructed to inspect systematic errors across test-retest sessions that depend on the range of each metric [54].

Systematic learning effects within and across testing sessions were identified. This is important to distinguish between task-related motor learning and behavioural recovery when using the VPIT to analyze the effect of interventions. In more detail, metrics were visualized for each of the five repetitions at test and retest. In addition, the slope (*η*) between test and retest for the median across all five repetitions was estimated and normalized with respect to the range of observed values. Strong learning effects were present if a paired t-test indicated significant di erences between test and retest and was below or equal −6.35 [33].

Lastly, the correlations between conventional impairment scales and the VPIT metrics were calculated, for the most a ected body side (*ρ_ma_*), to advance the pathophysiological interpretation of the digital health metrics.

## 3 Results

Out of the 30 post-stroke subjects, the VPIT protocol on the first testing day was completed by 23 and 27 individuals with the most a ected and less a ected body side, respectively. The reason for subjects not completing the protocol were: inability to understand the task (1 subject), severe visual deficits (1 subject), severe sensorimotor impairments (less a ected side: 1 subject; most a ected side: 5 subjects). The age of the included subjects was 59 [40, 53, 69, 88] years (median [minimum, 25*^th^*-percentile, 75*^th^*-percentile, maximum]) with 14 of them being female. FMA-UE scores for the most a ected and less a ected sides were 49 [32, 40, 57, 61] and 65 [56, 63, 66, 66], respectively. ARAT scores for the most a ected and less a ected sides were 47 [30, 39, 55, 57] and 57 [45, 57, 57, 57], respectively. Detailed subject characteristics can be found in Table SM4.

Twenty-one subjects also participated in the retest protocol, with 18 and 21 successfully completing it with the most a ected and less a ected side, respectively. The time between test- and retest was 7.88 [2.86, 5.22, 16.13, 46.96] days. The time to administer the VPIT protocol (instructions, familiarization, and five repetitions) was 16.66 [8.95, 12.34, 26.04, 37.84] min and 9.99 [6.27, 7.85, 16, 37.46] min for the most a ected and less a ected side, respectively, during the first testing session.

### 3.1 Characterization of sensorimotor impairments and activity limitations

The presence of sensorimotor impairments and activity limitations on a population level can be found in Table 1. According to the defined criteria, the percentage of subjects with sensorimotor impairments on the most a ected and less a ected sides varied between 70.0%-100.0% and 9.1%-50.0%, respectively, depending on the conventional scale. Similarly, the percentage with activity limitations ranged from 65.0%-90.0% and 4.5%-54.5% for the most a ected and less a ected side, respectively. Depending on the metric, the VPIT indicated sensorimotor impairments in 10.0%-50.0% and 0.0%-31.8% of all individuals with the most a ected and less a ected side, respectively. In total, 90% and 50% of all individuals showed impairment in at least one VPIT metric with the most a ected and less a ected side, respectively.

**Table 1:** Characterization of impairments and activity limitations. Conventional assessments and the VPIT were used to define the presence of sensorimotor impairments and activity limitations. For the VPIT, NHPT, and BBT, abnormal behaviour was defined if task performance was outside the 95*^th^*-percentile of a normative reference population. According to the ARAT, activity limitations were present if the score was below 55. All other conventional scales indicated the presence of impairments if the full score was not reached. Only participants with all conventional scales available were used. In total, 90% and 50% of all individuals showed impairment in at least one VPIT metric with the most a ected and less a ected side, respectively. MAS: Modified Ashworth Scale; NHPT: Nine Hole Peg Test; EmNSA: Erasmus modifications to the Nottingham Sensory Assessment; BBT: Box and Block Test; ARAT: Action Research Arm Test; FMA-UE: Fugl-Meyer Assessment Upper Extremity.

|  | Percentage of subjects with disability |  |
| --- | --- | --- |
|  | Most affected side<br>n = 20 | Less affected side<br>n = 22 |
| **Conventional scales: impairments** |  |  |
| FMA-UE | 100.0% | 50.0% |
| MAS | 75.0% | 9.1% |
| EmNSA | 70.0% | 18.2% |
| **Conventional scales: activity** |  |  |
| BBT | 90.0% | 54.5% |
| ARAT | 70.0% | 4.5% |
| NHPT | 70.0% | 9.1% |
| **VPIT: impairments in activity context** |  |  |
| Log jerk transport | 45.0% | 8.2% |
| Log jerk return | 35.0% | 9.1% |
| SPARC return | 30.0% | 9.1% |
| Path length ratio transport | 45.0% | 4.5% |
| Path length ratio return | 35.0% | 13.6% |
| Velocity max. return | 50.0% | 31.8% |
| Jerk peg approach | 30.0% | 0.0% |
| Grip force rate num. peaks transport | 50.0% | 22.7% |
| Grip force rate SPARC transport | 10.0% | 9.1% |
| Grip force rate SPARC hole approach | 45.0% | 4.5% |

Examples for the relationship between the VPIT metrics and conventional scales are visualized in Figure 1 (all correlations in Table 2, confidence intervals in Table SM5). The following correlations were significant after Bonferroni correction: force rate SPARC transport with MOCA (*ρ_ma_*=-0.61**); jerk peg approach with BBT (*ρ_ma_*=-0.73**), ARAT (*ρ_ma_*=-0.65**), and NHPT (*ρ_ma_*= 0.64**). Further, the correlations of the following conventional scales of impairments with the activity domain were significant after Bonferroni correction: FMA-UE with BT (*ρ_ma_*= 0.66**); MAS with BBT (*ρ_ma_*=-0.65**); FMA-UE with ARAT (*ρ_ma_*= 0.82**); MAS with ARAT (*ρ_ma_*=-0.62**).

**Figure 1:**
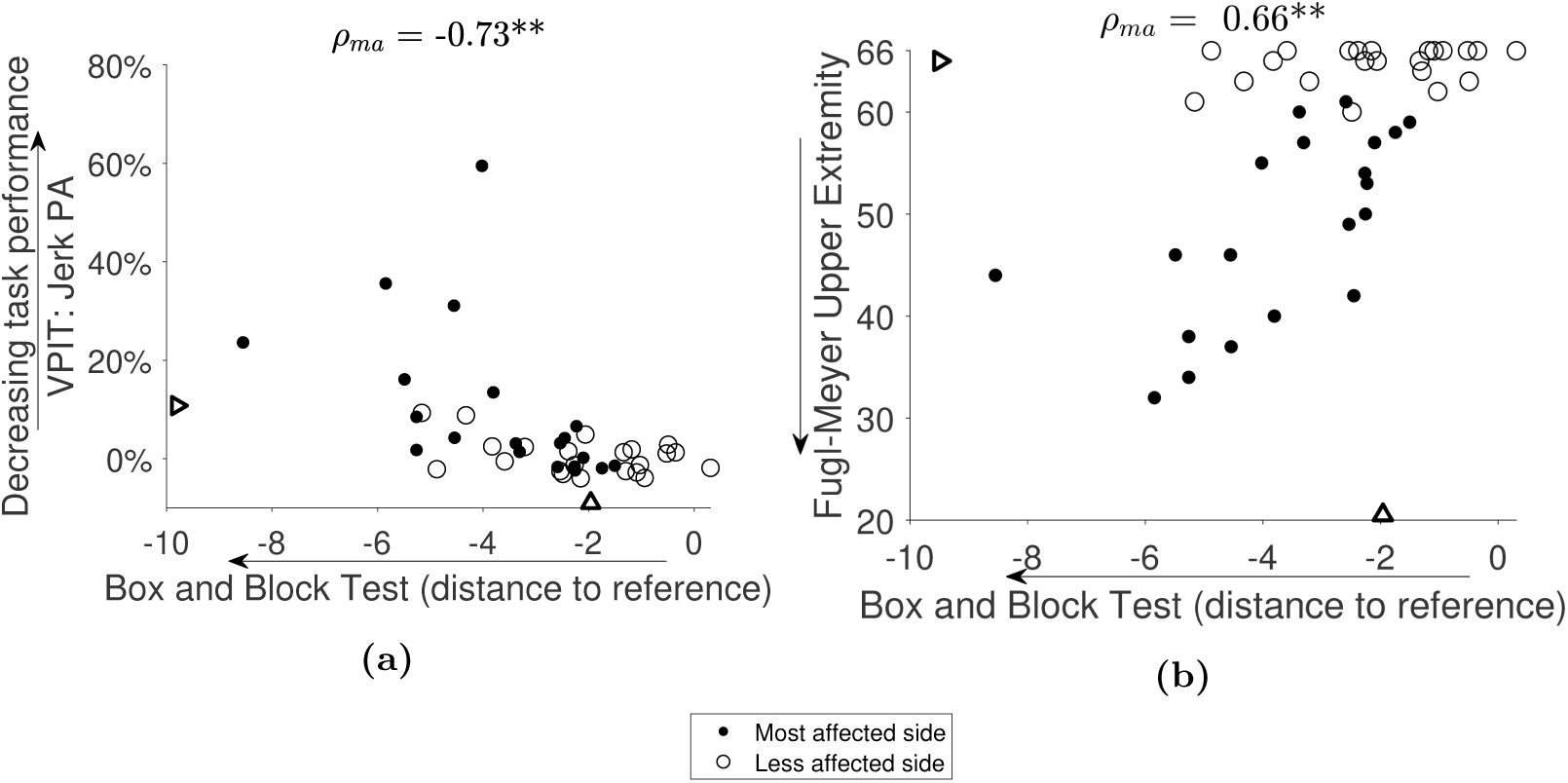
Example correlations between impairments (VPIT, Fugl-Meyer Upper Extremity) and activity limitations (Box and Block Test). The relationship of impairments and activity limitations was analyzed with Spearman correlations (*ρ*). Two pairs (a-b) were chosen for visualization purposes (all results in Table 2). Only data from the most a ected side (*ρ_ma_*) and the first testing session was used for the correlation analysis. For both VPIT and conventional scales, triangles represent a cut-o s indicating the presence of sensorimotor impairments (VPIT, Fugl-Meyer Upper Extremity) and activity limitations (Box and Block Test). A slightly stronger relationship was observed between impairments and activity limitations for the VPIT metric than the Fugl-Meyer assessment. **indicates *p*-value below the Bonferonni corrected significance level. VPIT: Virtual Peg Insertion Test.

**Table 2:**
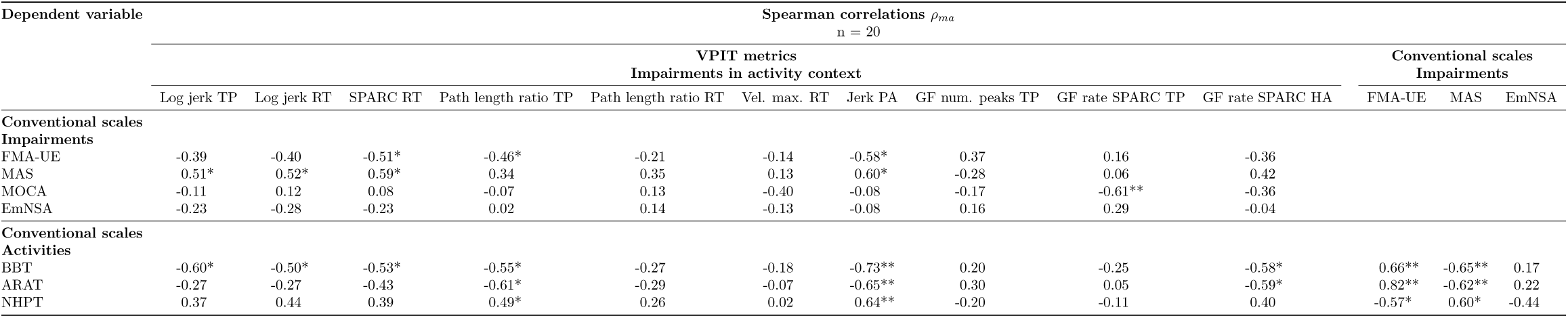
Correlation between conventional scales and VPIT metrics for the most a_ected side. Spearman correlation analysis was applied to analyze the relationship of conventional scales and VPIT metrics. Only data collected during the firrst testing session with the most affected body side was considered for this analysis. *indicates a p-value below 0.05 and **indicates a p-value below the Bonferroni-corrected signi_cance level. Bonferonni correction was applied within each table row. MAS: Modified Ashworth Scale; MOCA: Montreal cognitive assessment; NHPT: Nine Hole Peg Test; EmNSA: Erasmus MC modifications to the Nottingham Sensory Assessment; BBT: Box and Block Test; ARAT: Action Research Arm Test; FMA-UE: Fugl-Meyer Assessment Upper Extremity; GF: grip force. SPARC: spectral arc length. num: number. vel: velocity. TP: transport. RT: return. PA: peg approach. HA: hole approach. The Bonferroni-corrected signi_cance level was 0.05/13=0.0038 for the correlations with the BBT, ARAT, and NHPT, and 0.05/10=0.005 for all other conventional scales.

### 3.2 Test-retest reliability, measurement error, and learning effects of the VPIT metrics

Example visualization of the analyzed clinimetric properties can be found in Figure 2 (all metrics in Figure SM3, SM4, and SM7). The test-retest reliability and measurement error of all metrics are summarized in Table 3. The metrics fullfilling all criteria for the quality of the clinimetric properties were the *log jerk transport* (ICC 0.89, SRD% 23.31, −1.65), *log jerk return* (ICC 0.84, SRD% 28.56, η −4.85) and *force rate SPARC transport* (ICC 0.90, SRD% 20.49, η −5.02).

**Figure 2:**
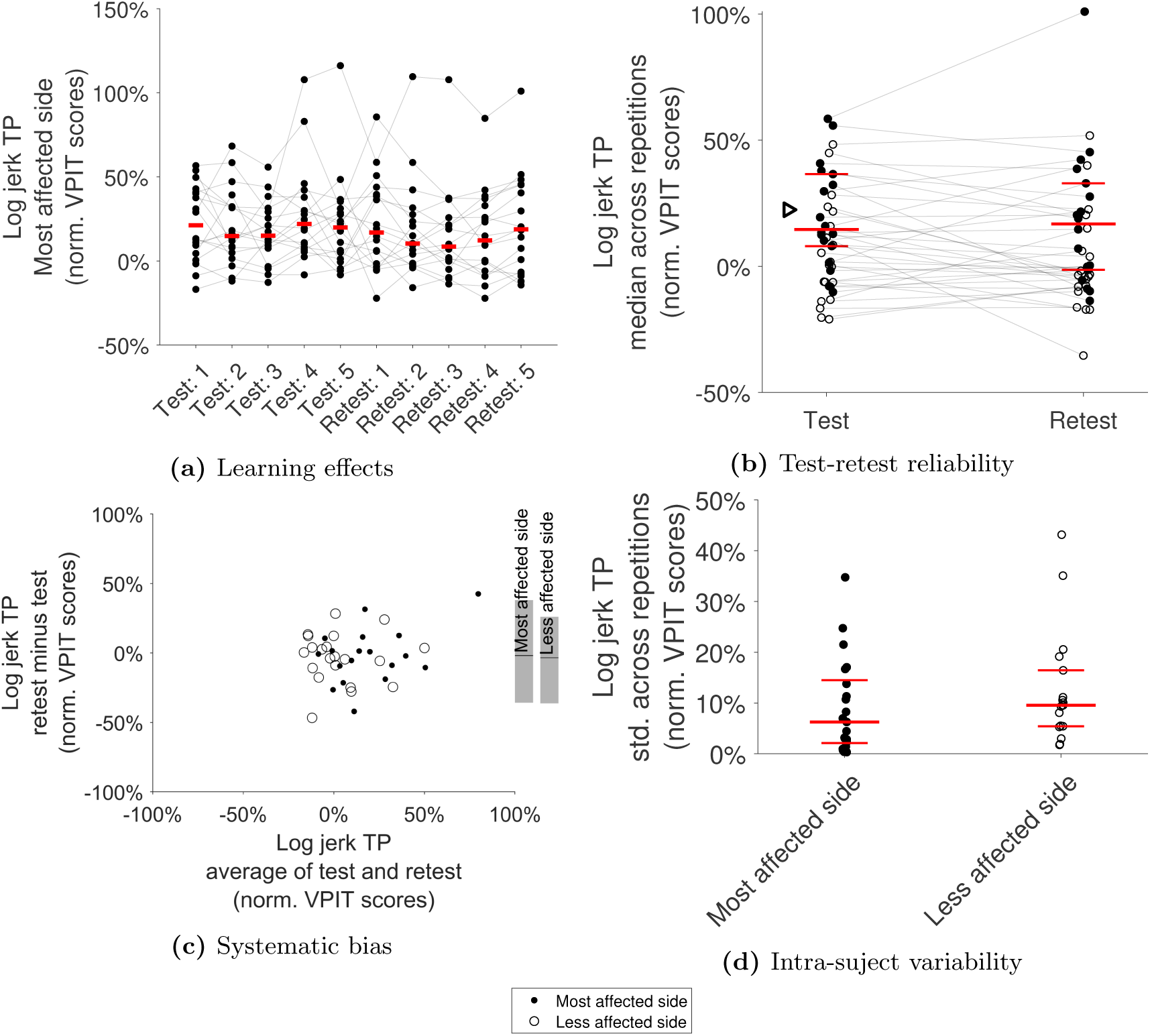
Clinimetric evaluation of the VPIT metrics: example log jerk transport. a) shows the behaviour of all subjects across five repetitions of test and retest to visualize potential learning effects. b) informs on test-retest reliability by visualizing the median across those five repetitions for test and retest. The red line indicates the population median for the most a ected side, the triangle corresponds to the 95*^th^*-percentile of the normative reference population, and shaded gray lines connect data from one subject. c) systematic bias was evaluated using a Bland-Altman plot (start and end of gray bars on the right indicate the 5*^th^*- and 95*^th^*-percentile). d) intra-subject variability was displayed through the standard deviation (std) within all ten repetitions of each subject. The example metric *log jerk transport* did not show strong learning effects, had high test-retest reliability, no systematic bias, and low intra-subject variability, therefore being defined as robust. TP: transport.

**Table 3:** Test-retest reliability: intra-class correlation (ICC) coe cients and smallest real di erences (SRD). The ICC (optimum at 1) describes the ability of a metric to discriminate between subjects across measurement sessions. The SRD% (optimum at 0%) describes a range of values for that the assessment cannot distinguish between measurement noise and an actual change in the underlying physiological construct. Bold ICC values represent acceptable test-retest reliability (i.e., above or equal 0.7). Bold SRD% indicate least strong measurement error (SRD%*<*30.3).

| Sensor-based metric | Test-retest reliability |  |  |  |
| --- | --- | --- | --- | --- |
|  | Most affected side<br>n = 18 |  | Less affected side<br>n = 21 |  |
|  | ICC [CI] | SRD% | ICC [CI] | SRD% |
| Log jerk transport | **0.89** [0.83, 0.92] | **23.31** | **0.79** [0.69, 0.86] | 30.79 |
| Log jerk return | **0.84** [0.75, 0.89] | **28.55** | **0.89** [0.84, 0.93] | **25.31** |
| SPARC return | **0.81** [0.72, 0.88] | 34.70 | **0.87** [0.81, 0.91] | **27.91** |
| Path length ratio transport | 0.58 [0.36, 0.72] | 54.05 | 0.66 [0.50, 0.77] | 52.38 |
| Path length ratio return | 0.49 [0.24, 0.66] | 52.24 | **0.84** [0.76, 0.89] | **29.09** |
| Velocity max return | **0.95** [0.92, 0.97] | **16.88** | **0.97** [0.93, 0.98] | **13.05** |
| Jerk peg approach | 0.48 [0.22, 0.65] | 94.55 | **0.92** [0.88, 0.95] | **19.75** |
| Grip force rate num. peaks transport | **0.87** [0.80, 0.91] | **24.58** | **0.90** [0.84, 0.93] | **21.70** |
| Grip force rate SPARC transport | **0.90** [0.85, 0.94] | **20.49** | **0.89** [0.84, 0.93] | **21.43** |
| Grip force rate SPARC hole approach | **0.85** [0.72, 0.91] | 34.20 | **0.78** [0.67, 0.85] | 41.39 |

The metrics having insu cient (ICC*<*0.7) test-retest reliability were *path length ratio transport/return* and *jerk peg approach* for the most a ected side and *path length ratio transport* for the less a ected side. Systematic bias across test-retest session according to Bland-Altman plots was visible especially for *path length ratio transport/return* and *jerk peg approach*. The metrics *SPARC return*, *path length ratio transport/return*, *jerk peg approach*, and *grip force rate SPARC hole approach* for the most a ected side as well as *log jerk transport*, *path length ratio transport*, and *grip force rate SPARC hole approach* for the less a ected side did not pass the measurement error evaluation (SRD%*>*30.3).

On the most a ected side, learning effects across test-retest were strong (*p*-value*<*0.05 and *η >-*6.35) for *path length ratio transport*, *velocity max. return*, *force rate num. peaks transport*, and *force rate SPARC hole approach* (Table SM6, Figure SM5). For the less a ected side, learning effects were strong for *velocity max. return* and *force rate num. peaks transport* (Table SM6, Figure SM6).

## 4 Discussion

The aim of this work was to evaluate the potential of digital health metrics, extracted from a technology-aided assessment (VPIT), in chronic post-stroke individuals, to inform on arm and hand sensorimotor impairments and especially characterize their functional relevance. In addition, the objective was to establish the interpretation and robustness of the metrics in this population to pave the way for their integration into clinical trials. The novelty of this work lies in the application and evaluation of a technology-aided assessment that has high clinical applicability and allows rapidly capturing movement and grip force patterns during a goal-directed, functionally relevant manipulation task without providing arm weight support. Hence, we expected that the metrics provide a fine-grained, robust, and clinically applicable assessment of sensorimotor impairments in arm and hand with functional relevance. This hypothesis was evaluated in 30 chronic post-stroke subjects. Twenty-three of these, the ones with mild to moderate sensorimotor impairments and without strong cognitive deficits, were able to successfully complete the goal-directed manipulation task protocol with their most a ected body side, thereby confirming previous reports about the feasibility of such tasks in individuals with mild to moderate neurological deficits [30, 32, 55].

### 4.1 Assessment of functionally relevant sensorimotor impairments with a technology-aided goal-directed manipulation task

The digital health metrics allowed identifying a high amount of individuals with impairments in the most a ected (90%) and less a ected (50%) side. This could only be achieved by considering multiple kinematic and kinetic metrics, thereby providing the envisioned fine-grained assessment of arm and hand sensorimotor deficits. Nevertheless, conventional assessments detected sensorimotor impairments (FMA-UE 100% for most a ected side) in more post-stroke individuals than the digital health metrics, even though the latter have a more sensitive scale without ceiling effects. We argue that the reduced rate of detected impairments with the digital health metrics is because individuals can compensate for certain impairments through the redundancy of the human motor apparatus and therefore still achieve normal performance during the goal-directed tasks [13, 38, 56].

Moreover, the digital health metrics showed high significant correlations with the BBT and moderate significant correlations with the ARAT and NHPT. This suggests that the goal-directed manipulation task is able to describe sensorimotor impairments that are functionally relevant and especially related to the ability to repeatedly grasp and transport lightweight objects as well as dexterous finger manipulations. Indeed, it is intuitive that the goal-directed manipulation task is especially related to the BBT, given the similar movements that are required to complete the two tests. In addition, the correlations of the digital health metrics with the BBT and NHPT were slightly higher than the ones observed between conventional assessment of sensorimotor impairments (FMA-UE, MAS, EmNSA) and BBT and NHPT. We speculate that this slightly stronger relationship results from the digital health metrics being recorded during a functional task, whereas conventional assessments of impairments describe them in the absence of a functional context. For the ARAT, the correlations were considerably higher with the FMA-UE than with the digital health metrics. Compared to the technology-aided task, the FMA-UE and ARAT emphasize more the ability to flex the shoulder, thereby explaining their strong relationship that has also been extensively reported in literature [11–13].

When relating these insights to the state of the art, it becomes apparent that only few technology-aided approaches quantify movements without arm weight support and also include object manipulations with the hand, which are especially important to linking impairments and activity limitations [24–27]. For example, Alt Murphy et al. showed similar correlation, as reported herein, between movement smoothness and the ARAT for post-stroke subjects that performed a drinking task recorded with an optical motion capture system [24, 25]. Similarly, Johansson and Häger used an optical motion capture system for characterizing kinematics during a modified version of the NHPT and found high correlations between movement smoothness and the task completion time [27]. While these approaches are promising to relate sensorimotor impairments and activity limitations and further also allow to study compensatory trunk movements, the solutions rely on a costly and time-consuming measurement setup with an optical motion capture system, thereby having limited clinical applicability. Research towards more rapidly applicable approaches has also been proposed, for example relying on the same robotic end-e ector as the VPIT [57, 58]. However, the presented task did not require any precise object manipulations and relied on the regular handle of the end-e ector that cannot record grip forces. Unsurprisingly, the correlations with the activity domain were considerably lower (multiple regression R^2^ up to 13% for ARAT, which would correspond to a Pearson correlation of 36% for the univariate case). Lastly, it is important to emphasize that such approaches are especially tailored for individuals with mild to moderate neurological deficits, and diverging results can be observed in subjects with more severe impairments [59–62]. This stems from such individuals typically having only limited residual ability to use the hand, which makes the assessment of arm impairments su cient to establish a link between impairments and activity limitations. Also, severely impaired individuals typically require arm weight support, thereby shadowing the influence of functionally relevant impairments such as weakness [19].

Hence, the proposed technology-aided assessment crystallizes as an interesting solution allowing a rapid (median 16.6 min with most a ected side including instructions) and, relative to optical motion capture systems or exoskeletons, inexpensive (approx. 4000 USD hardware costs) assessment of sensorimotor impairments in arm and hand in individuals with mild to moderate disability. Moreover, the impairments detected with the technology-aided approach showed relevance for performing activities similar to the NHPT and BBT, which was enabled by the task involving precise manipulations, the absence of arm weight support, and the quantification of grip forces.

### 4.2 Pathophysiological correlates of VPIT metrics and functional relevance of impairments

While conventional assessments (FMA-UE, MAS, EmNSA) capture sensorimotor impairments without functional context, it was still expected to observe moderate correlations between functionally relevant impairments and VPIT metrics. These correlated with the MAS and FMA-UE, which suggests that the metrics are sensitive to increased muscle tone and abnormal coupling of the shoulder, arm, and hand. While trends were visible for many metrics, the strongest ones were found for the metric *jerk peg approach*, which was also correlated most strongly to conventional scales of activity. This metric describes especially the precise coordination of movements and the release of grip forces that is required to insert a peg, which might be modulated by the integrity of the corticospinal tract [33, 63]. This idea is supported by the correlation with the FMA-UE and MAS, given that the abnormal coupling of joints is expected to be driven by corticospinal tract integrity, which can also contribute to increased muscle tone, depending on lesion location and severity [64–67]. However, these speculative statements require further validation, given that the correlations with the FMAUE and MAS were not significant after Bonferroni correction, and that neurophysiological markers would be required for making strong conclusions. Also, a clear correlation of the FMA-UE with NHPT (not significant after Bonferroni), BBT, and ARAT was observed, which suggests either the functional relevance of the ability to perform fractionated movements with single joints, expected to be driven by corticospinal tract integrity, or the co-occurrence of other impairments when the main neural transmission pathway is disrupted. Given that subjects often perform compensatory movements allowing to improve task performance in the presence of abnormal joint couplings [13, 38], we speculate that the latter option is not unlikely.

Although the clinical importance of spasticity post-stroke is subject to critical discussions, the results indicating a reduced ability to perform goal-directed activities in individuals with increased muscle tone are in line with previous literature [68, 69].

Somatosensory impairments, as assessed by the EmNSA, were not significantly correlated to any VPIT metrics and did not contribute to functional task performance in the conventional scales. Interestingly though, moderate correlations (significant before Bonferroni) were found for the *force rate SPARC hole approach* metric and the BBT and ARAT. Given that this metric characterizes grip force coordination and is expected to be influenced by sensory deficits [33], we speculate that these deficits might have not been captured by the clinical scale of sensory impairments that is well known to lack sensitivity [70].

The only VPIT metric being significantly correlated to the MOCA as a general descriptor of cognitive impairments was the force rate SPARC transport. This might result from a misunderstanding of the visual feedback provided by the task and the subsequent uncoordinated application of grip forces. However, as only one metric was a ected, this also indicates only a minor influence of cognitive deficits on the perception of the virtual environment or understanding of the VPIT task.

These results showing moderate correlations between conventional impairment scales and digital health metrics are in general in line with literature, even though the relationships are strongly context-dependent [17, 71–73].

### 4.3 Clinimetric properties of the VPIT metrics

The clinimetric properties of the ten VPIT core metrics were previously positively evaluated in una ected subjects [33]. Also, a first preliminary evaluation of the VPIT was done in post-stroke subjects [32]. However, this evaluation relied on a di erent measurement protocol and did not yet consider the recently introduced ten core metrics, which were selected by applying conservative and objective selection criteria [33]. Herein, we confirm the robustness of three VPIT core metrics, *log jerk transport* (ICC 0.89, SRD% 23.31, *η* −1.65), *log jerk return* (ICC 0.84, SRD% 28.56,η −4.85), and *force rate SPARC transport* (ICC 0.90, SRD% 20.49, −5.02) in the most a ected side of chronic post-stroke subjects. This implies that these metrics are highly reliable, have no strong measurement error, and are not showing strong learning effects. This is expected to make the metrics suitable for assessing sensorimotor impairments in a longitudinal manner. Given the previous validation, all ten metrics can still be used to detect the presence of sensorimotor impairments in cross-sectional studies [33]. Reasons why the metrics were more robust in neurologically intact than a ected subjects might be the smaller sample size used for the analysis in this work as well as higher intra-subject variability in post-stroke subjects (Figure 2 and SM7). This rather high variability might be because the VPIT allows heterogeneous task completion strategies and the haptic device being able to render only up to 3.3 N of haptic feedback, which can lead to an unstable haptic rendering of the virtual reality environment. Also, the variability might be influenced by a visuomotor transformation from the end-e ector to the virtual reality environment that has to be learned throughout multiple repetitions of the task (Figure SM5), as also observed in other virtual reality-based assessments [74].

It is challenging to compare the clinimetric properties of the VPIT metrics to the ones extracted from other technology-aided assessments due to the context-dependence of metrics [17, 73]. Moreover, there is a lack of quality in the evaluation of technology-aided assessments and in-depth and thorough validation is only rarely implemented [17]. In the few cases where measurement error has been reported, its magnitude was again dependent on the assessment metric and platform, with overall mostly similar ranges (e.g., SRD of 13.2% to 95.0%) to the VPIT metrics [62,75–79]. Compared to conventional assessments (e.g., FMA-UE measurement error of 7.9%; ARAT of 6.1%) [76, 80], the measurement errors of most technology-aided assessment metrics seem consistently elevated, even though comparisons are also challenged by the use of di erent SRD implementations. Nevertheless, we argue that this results from technology-aided assessments providing a fine-grained picture of the behavioural components underlying task performance, which makes them more susceptible to behavioural variability compared to the often ordinal outcome measures of conventional scales. Hence, we recommend researchers to thoroughly evaluate the clinimetric properties of technology-aided assessments and especially consider intra-subject variability as an important factor when designing assessment tasks. This is fundamental to fulfil the high expectations of the research community about technology-aided assessments providing more sensitive outcome measures than conventional scales.

### 4.4 Limitations

The major limitation of this work is the rather small amount of post-stroke participants included in the analysis, which limits the generalizability of the results to other individuals that potentially show di erent impairment phenotypes. This also led to rather high confidence intervals (Table SM5) for the correlation analysis and emphasizes the need for further validation. Further, compensatory movements, for example by the trunk, were not captured by the end-e ector based approach, but might be important to fully understand the relationship between impairments and activity limitations.

## 5 Conclusions

This work provides evidence about the importance of technology-aided assessments that are considering precise goal-directed manipulations and grip forces without arm weight support, such as the VPIT. These approaches can enable a robust, sensitive, and objective way to assess arm and hand sensorimotor impairments that are functionally relevant in chronic post-stroke individuals with mild to moderate deficits. Further, the VPIT allowed implementing such an approach in a highly clinically applicable manner, by being rapidly applicable and, for a technology-aided assessment, inexpensive. This promises to better identifying impairments with high functional relevance as therapy targets in clinical research and practice, which might ultimately contribute to optimizing the functional benefits of neurorehabilitation interventions.

In the future, it should be explored whether the assessment with the VPIT provides clinical benefits when used as a complementary source of information in clinical practice. Further, the presented results should be confirmed within large-scale trials, where structural neuroimaging markers together with clustering approaches should be used to fully unravel the pathophysiological correlates of digital health metrics.

## Acknowledgements

The authors would like to thank Sascha Motazedi Tabrizi for support during data collection.

## Funding

This project received funding from the European Union’s Horizon 2020 research and innovation programme under grant agreement No. 688857 (SoftPro) and from the Swiss State Secretariat for Education, Research and Innovation (15.0283-1). The authors declare that the funding bodies did not influence the design of the study, the collection, analysis, and interpretation of data, and the writing of the manuscript.

## Ethical approvals

The study was registered at clinicaltrials.gov (NCT03135093) and approved by the responsible Ethical Committees (ID 2016-02075 and BASEC:2017-00398).

## Conflicts of interest

The authors declare no conflicts of interest.

## Data availability

The data presented in this manuscript are available upon reasonable request and under consideration of the ethical regulations.

## Author contributions

Study design: CK, AS, JH, AL, RG, OL. Data collection: CK, AS, JH. Data analysis: CK. Data interpretation: CK, RG, OL. Manuscript writing: CK, RG, OL. Manuscript review: CK, AS, JH, AL, RG, OL. All authors read and approved the final manuscript.

## List of abbreviations

ARAT: Action Research Arm Test. BBT: Box and Block Test. Erasmus modified Nottingham sensory assessment. FMA-UE: Fugl-Meyer Assessment Upper Extremity. GF: Grip Force. HA: Hole Approach. ICC: Intra-class Correlation Coe cient. MAS: Modified Ashworth Scale. MOCA: Montreal Cognitive Assessment. Num: Number. PA: Peg Approach. RT: Return. SRD: Smallest Real Di erence. SPARC: Spectral Arc Length. TP: Transport. Vel: Velocity. VPIT: Virtual Peg Insertion Test.

## Ethical approvals

The study was registered at clinicaltrials.gov (NCT02688231) and approved by the responsible Ethical Committees (University of Leuven, Hasselt University, and Mariaziekenhuis Noord-Limburg).

## Conflicts of interest

The authors declare no conflicts of interest.

## Consent for publication

Not applicable.

---

**Table SM4:**
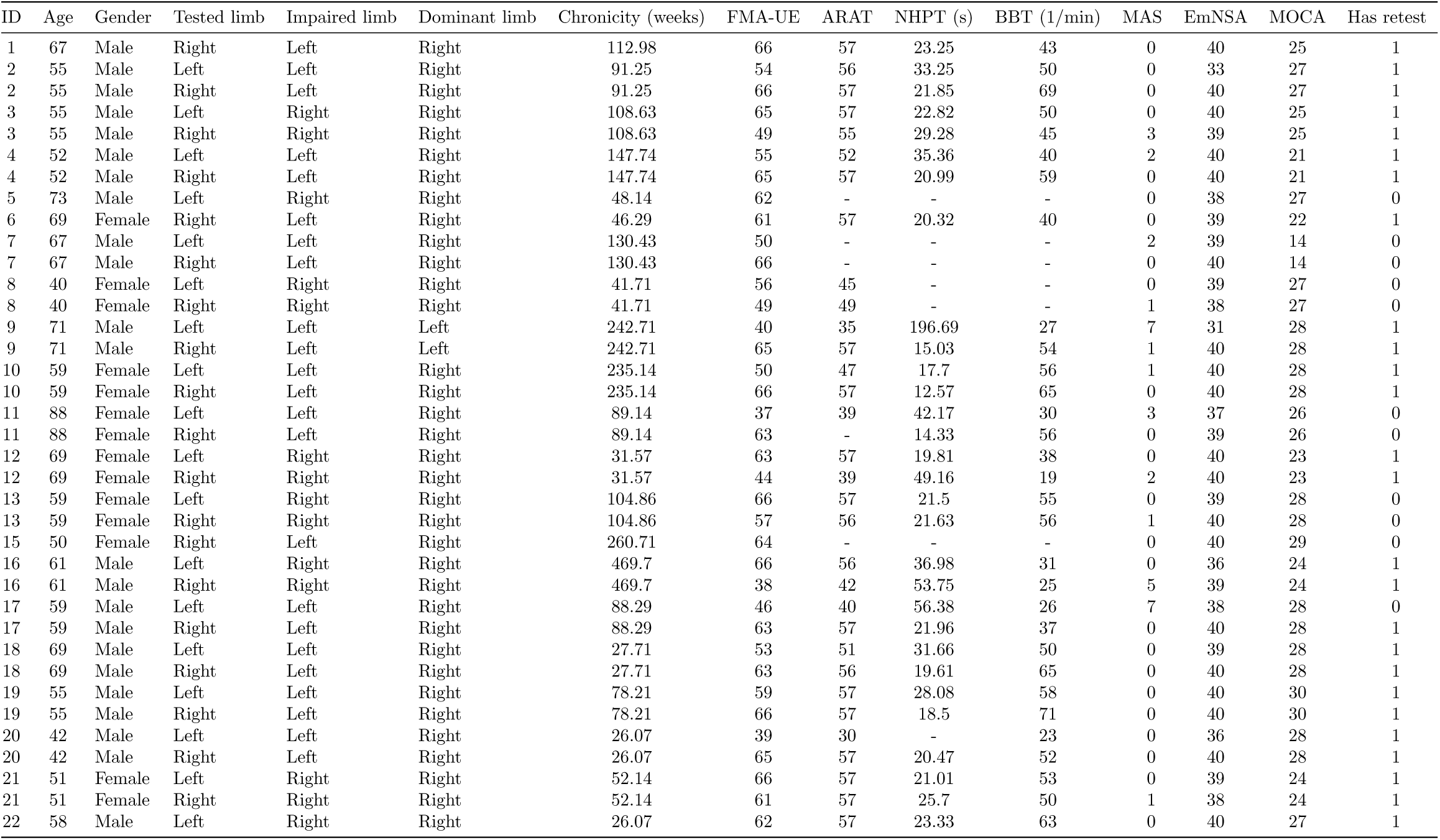
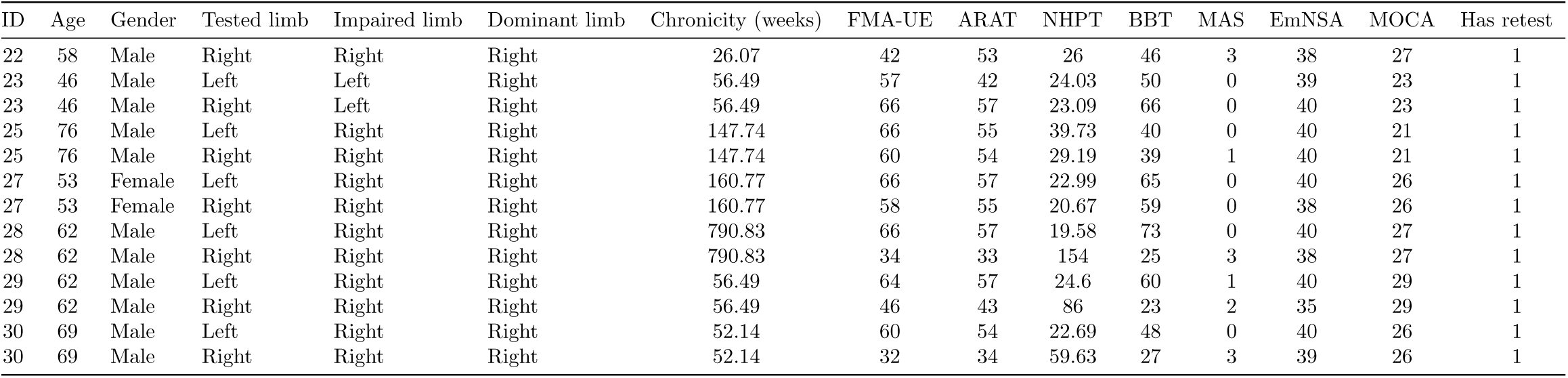
Demographic and clinical information for all post-stroke subjects. FMA-UE: Fugl-Meyer Assessment Upper Extremity. ARAT: Action Research Arm Test. NHPT: Nine Hole Peg Test. BBT: Box and Block Test. MAS: Modi_ed Ashworth Scale (sum across muscle groups). EmNSA: Erasmus modi_ed Nottingham Sensory Assessment. MOCA: Montreal Cognitive Assessment.

**Table SM5:**
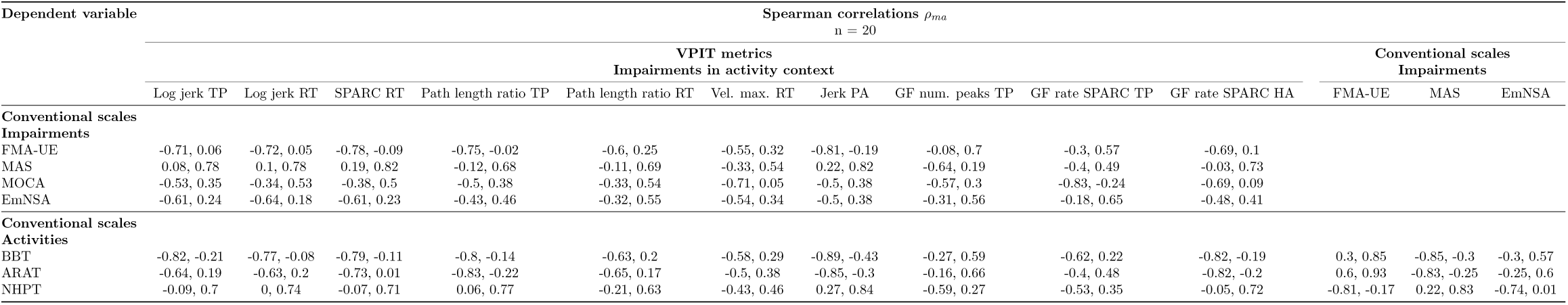
Confidence intervals for correlation between conventional scales and VPIT metrics for the most affected side. Spearman correlation analysis was applied to analyze the relationship of conventional scales and VPIT metrics. Ninety-five percent con_dence intervals were constructed via Fisher’s z-transform and reported as ‘lower bound, upper bound’. Only data collected during the firrst testing session with the most affected body side was considered for this analysis.MAS: Modified Ashworth Scale; MOCA: Montreal cognitive assessment; NHPT: Nine Hole Peg Test; EmNSA: Erasmus MC modifications to the Nottingham Sensory Assessment; BBT: Box and Block Test; ARAT: Action Research Arm Test; FMA-UE: Fugl-Meyer Assessment Upper Extremity; GF: grip force. SPARC: spectral arc length. num: number. vel: velocity. TP: transport. RT: return. PA: peg approach. HA: hole approach.

**Table SM6:** Quantification of learning effects. A linear regression model was used to test, per metric, whether a systematic improvement, indicative of learning effects, between test and retest was present. The slope *η* was normalized relative to the range of observed values. Bold entries indicate metrics without strong learning effects (e ect non-significant or *η >*-6.35).

| Sensor-based metric | Learning effects ($\eta$) | |
| --- | --- | --- |
|  | Most affected side<br>n = 18 | Less affected side<br>n = 21 |
| Log jerk transport | **-1.65** | **-4.00** |
| Log jerk return | **-4.85** | **-6.67** |
| SPARC return | **-8.10** | **-6.67** |
| Path length ratio transport | **-7.68** | **0.80** |
| Path length ratio return | -13.28 | **-4.23** |
| Velocity max. return | -8.86 | -8.90 |
| Jerk peg approach | **-9.92** | **3.21** |
| Grip force rate num. peaks transport | -10.16 | -7.16 |
| Grip force rate SPARC transport | **-5.02** | **-4.64** |
| Grip force rate SPARC hole approach | -18.90 | **-5.46** |

**Figure SM3:**
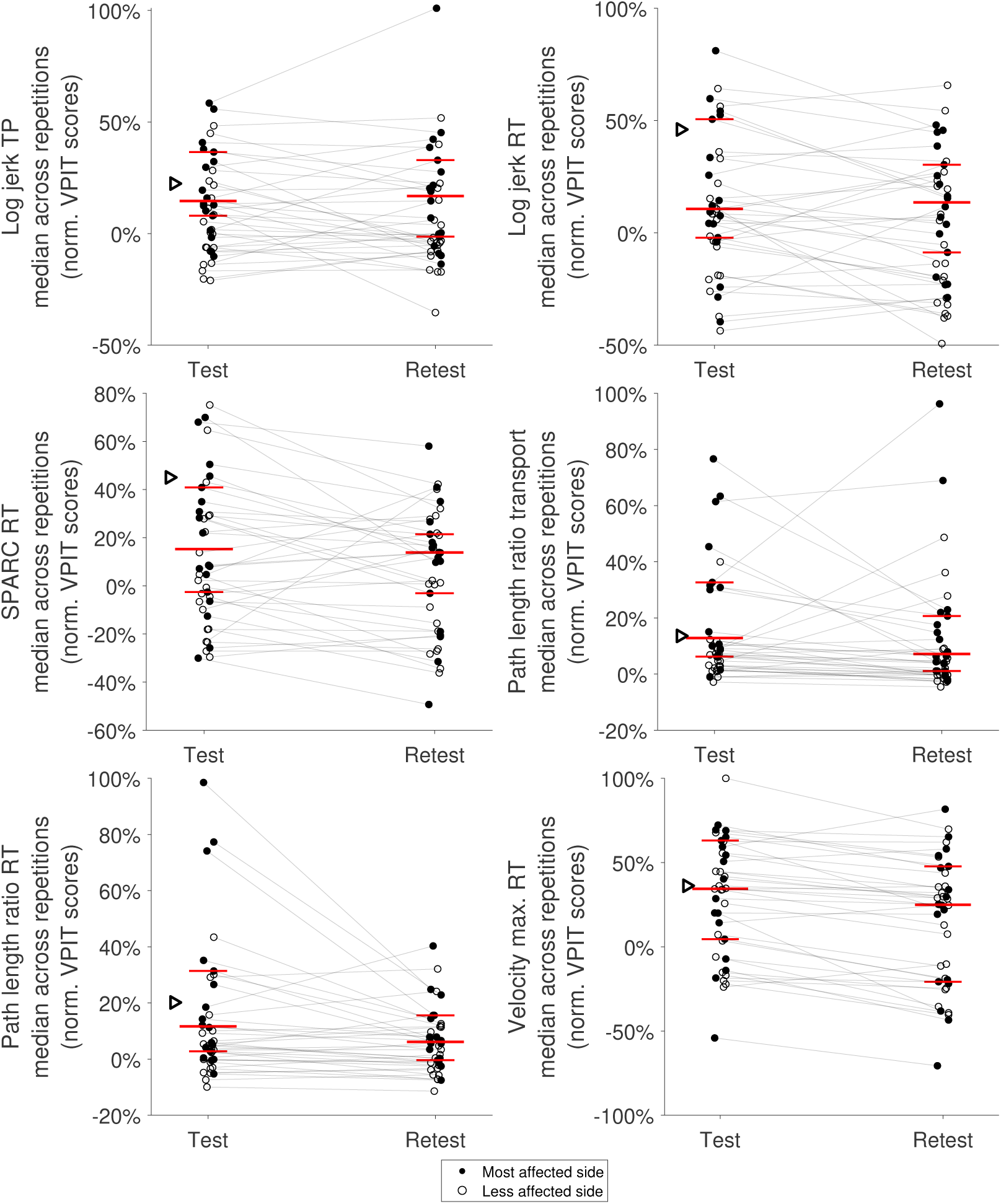
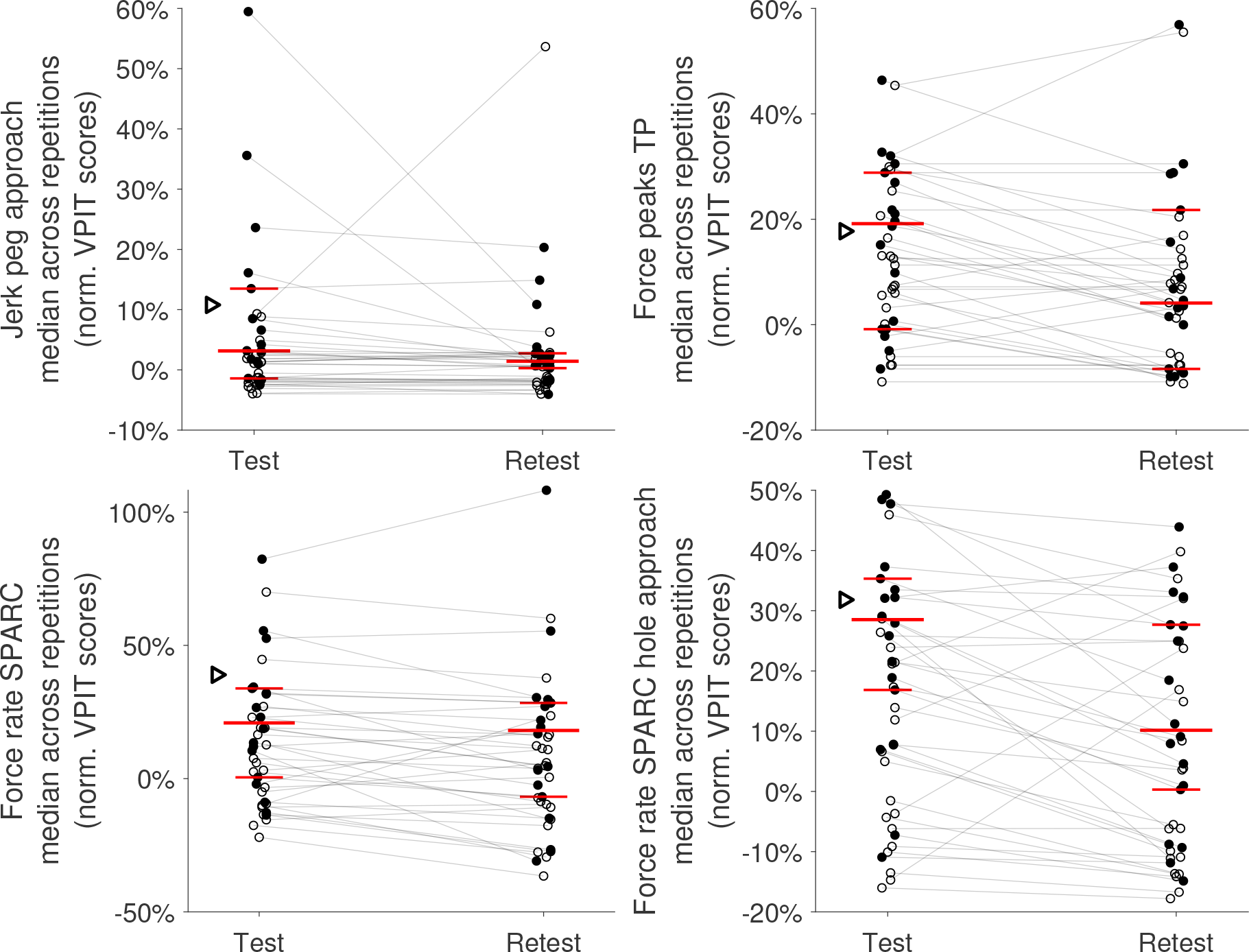
Test and retest scores for all VPIT metrics. The task performance level decreases with increasing VPIT scores. Additionally, 0% represents the median of an unimpaired reference population and 100% the task performance of the worst neurological subject in the VPIT database. The long red horizontal bar indicates the population median for the most a ected side. The shorter red horizontal bars represent the 25*^th^*- and 75*^th^*-percentile. The black triangle represents the 95*^th^*-percentile of the unimpaired reference population. Pre- and post-measurements of a single subject and body side are connected with a gray line.

**Figure SM4:**
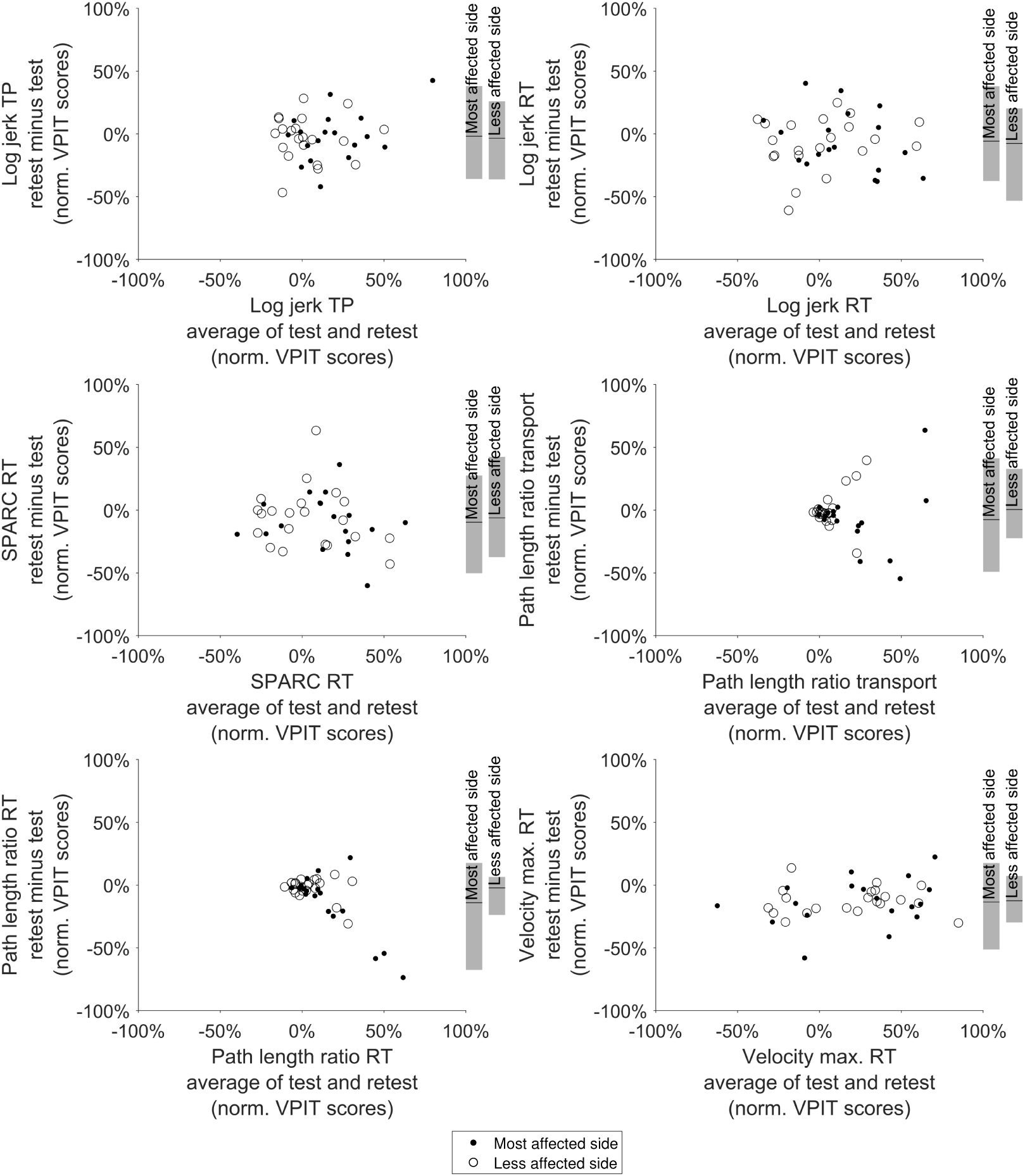
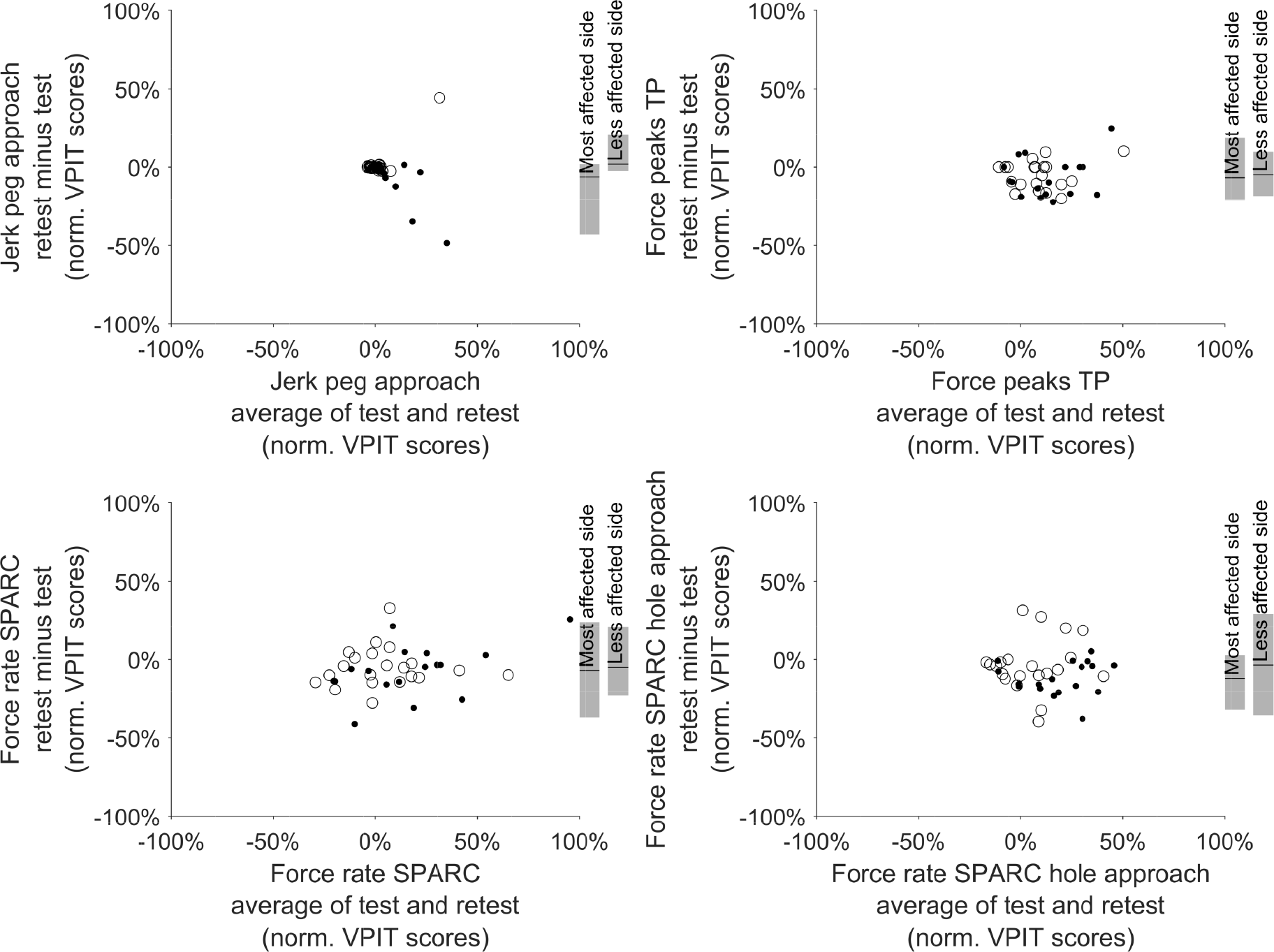
Bland-Altman plots for the sensor-based metrics of the VPIT. The vertical axis represents the di erence between the test and retest measurement, whereas the horizontal axis represents their average. The dashed horizontal line represents ideal behaviour (i.e., zero di erence between measurements). Further, the solid black horizontal bars and the shaded gray areas represent the median, 5*^th^* and 95*^th^*-percentile, respectively, for subjects tested on the most and less a ected side. TP: transport. RT: return.

**Figure SM5:**
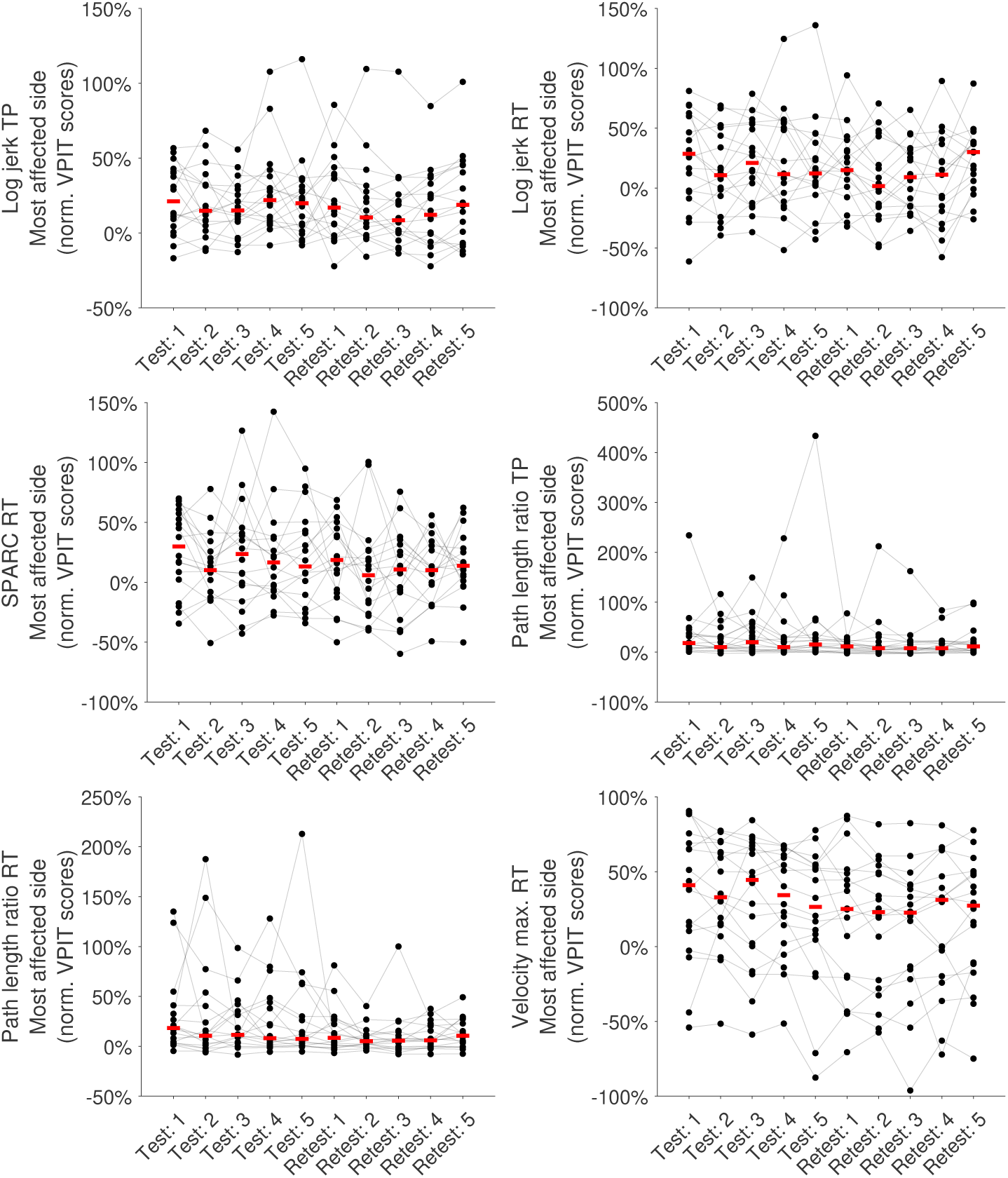
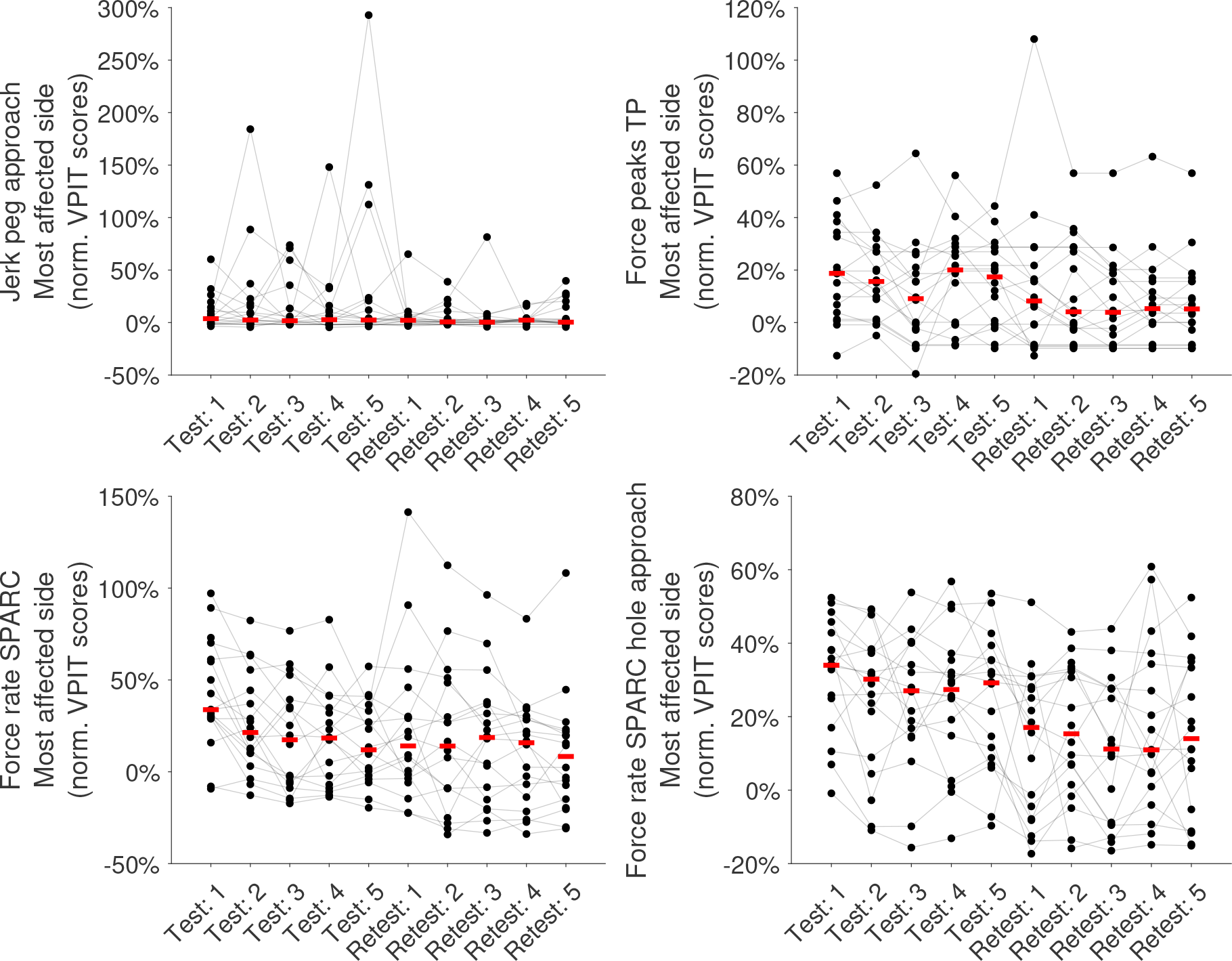
Learning effects in the VPIT metrics for the most affected side. The behaviour of all subjects across five repetitions of test and retest is visualized to identify potential learning effects. Gray horizontal lines connect the data of one individual. The red line indicates the median across subjects. TP: transport. RT: return. SPARC: spectral arc length.

**Figure SM6:**
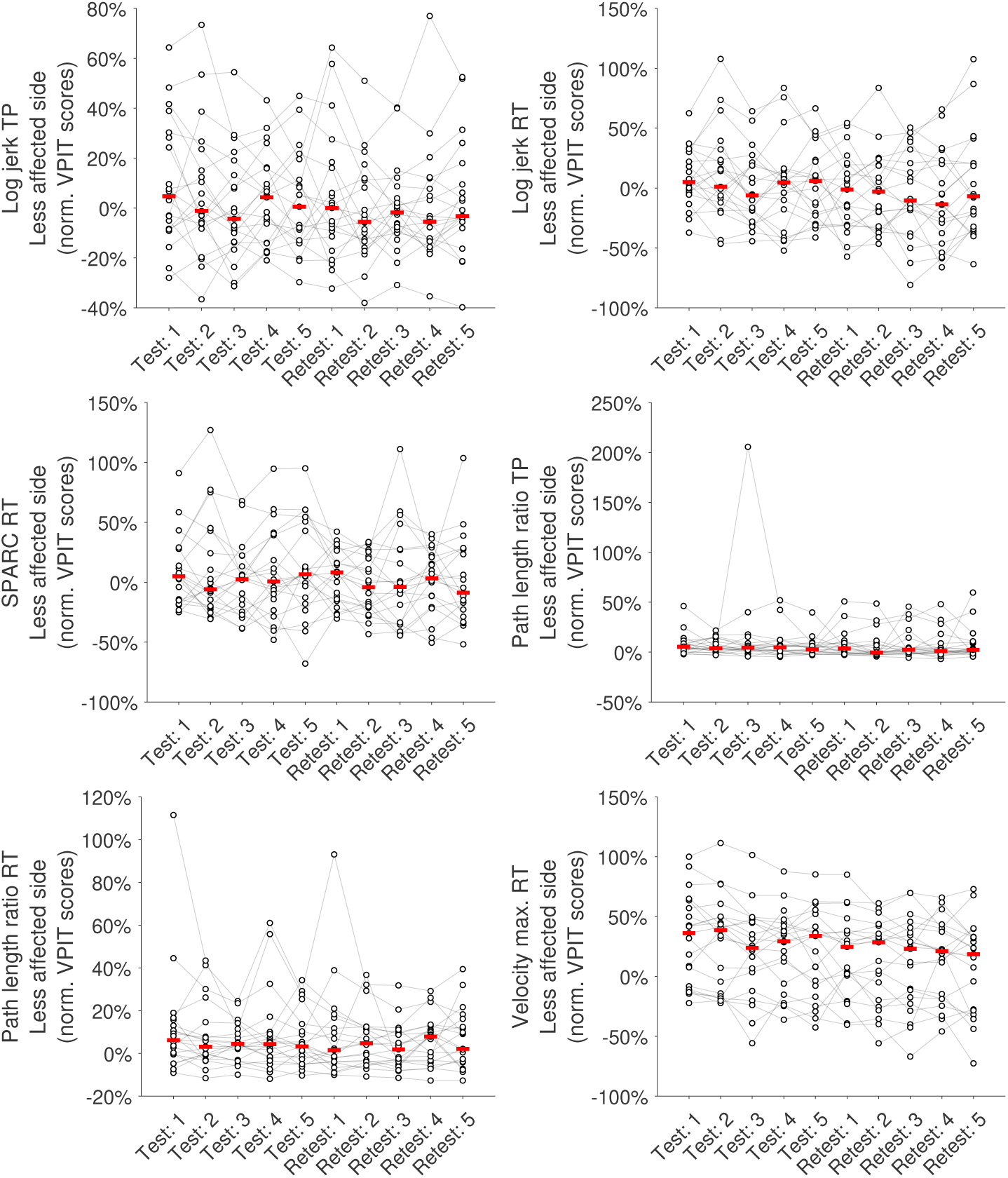
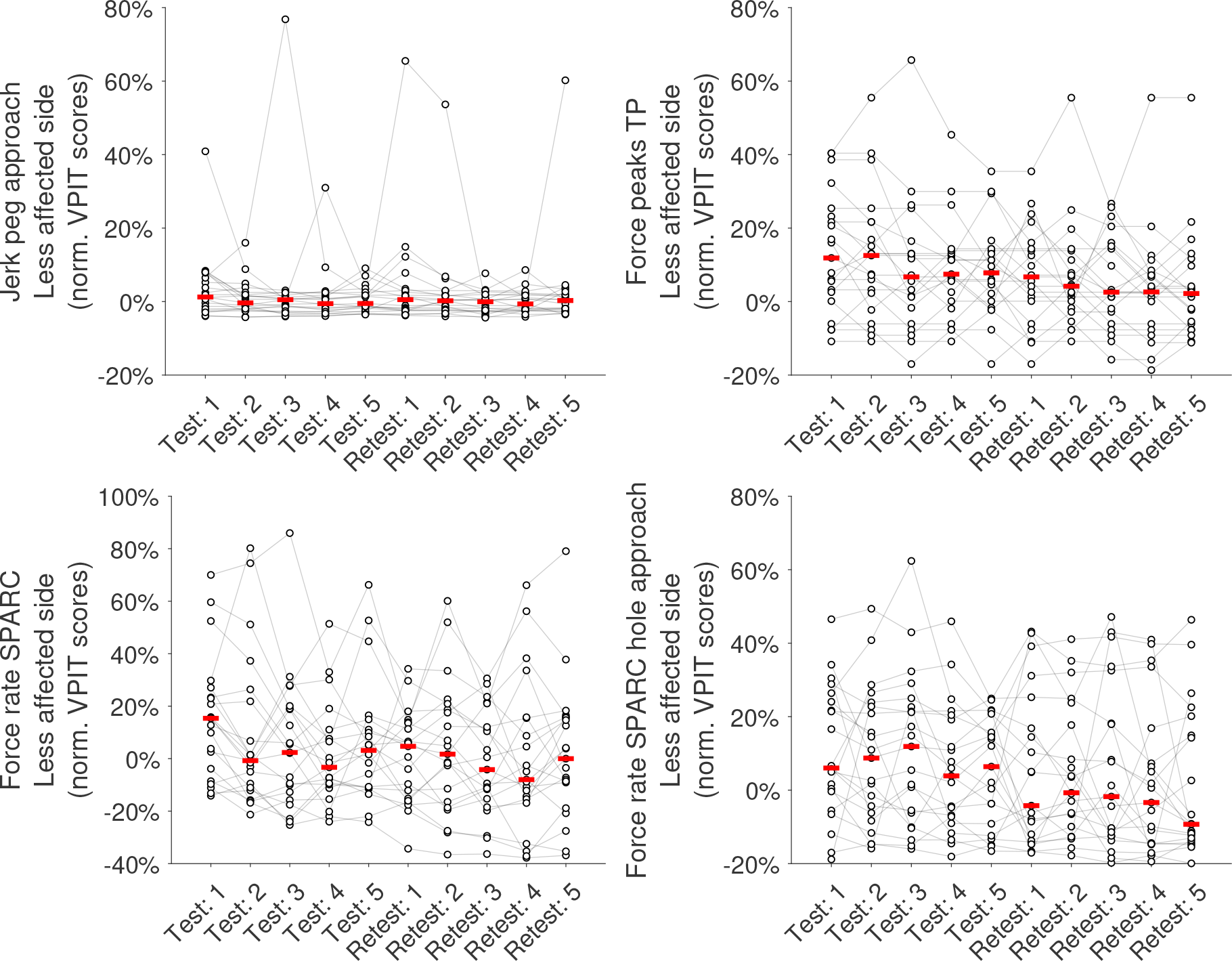
Learning effects in the VPIT metrics for the less a ected side. The behaviour of all subjects across five repetitions of test and retest is visualized to identify potential learning effects. Gray horizontal lines connect the data of one individual. The red line indicates the median across subjects. TP: transport. RT: return. SPARC: spectral arc length.

**Figure SM7:**
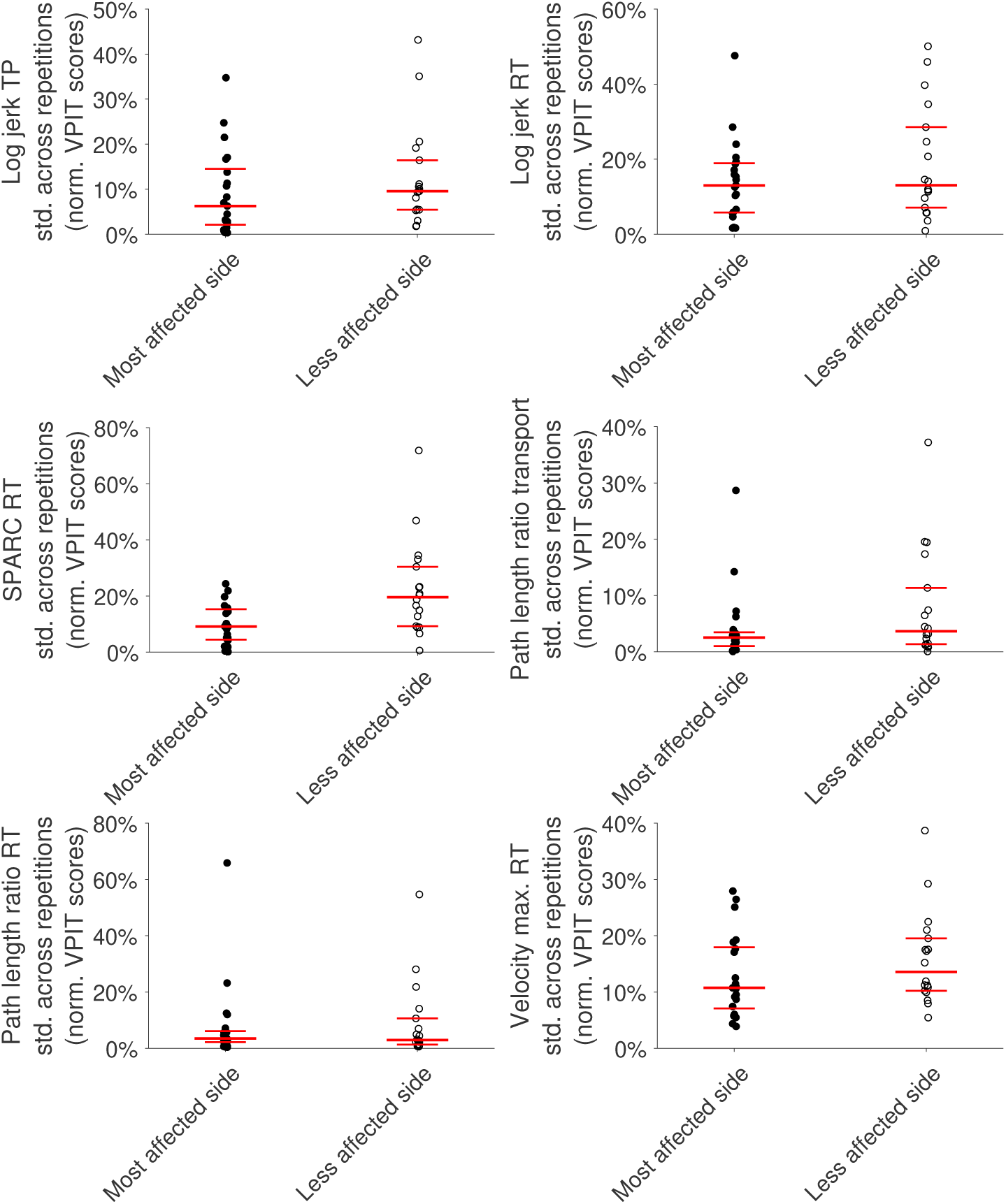
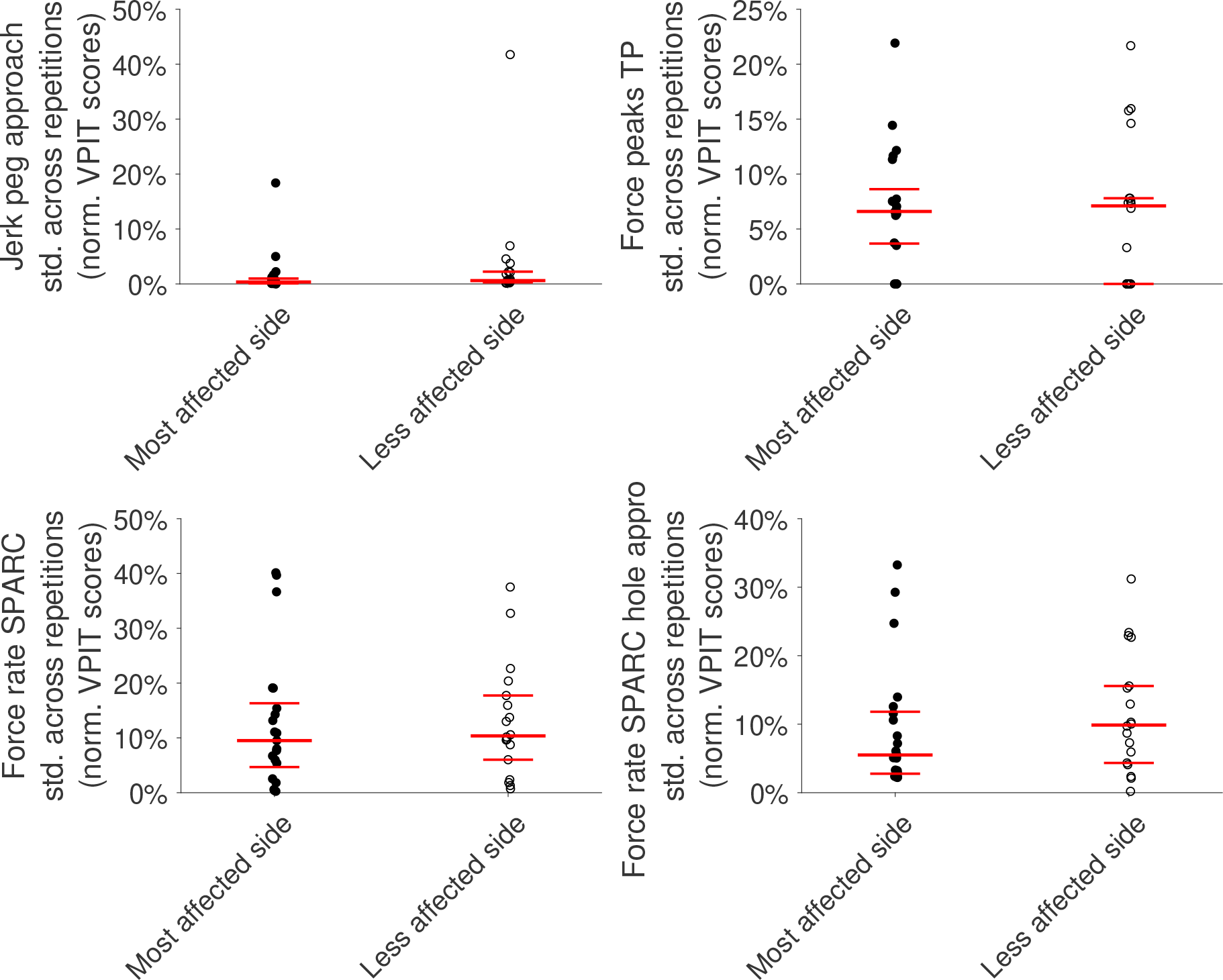
Intra-subject variability of the VPIT metrics. The standard deviation within the ten repetitions of the VPIT of each subjects was visualized. The longest red line indicates the population median, whereas the shorter red lines indicate the 25*^th^* and 75*^th^*-percentiles. TP: transport. RT: return. SPARC: spectral arc length.

## Notes

### Competing Interest Statement

The authors have declared no competing interest.

